# Evolutionary analysis of V protein pseudogenization in an RNA editing-deficient paramyxovirus

**DOI:** 10.64898/2026.04.06.716634

**Authors:** Tofazzal Md Rakib, Lipi Akter, Yusuke Matsumoto

**Affiliations:** Transboundary Animal Diseases Research Center, Joint Faculty of Veterinary Medicine, Kagoshima University, Kagoshima 890-0065, Japan; Department of Pathology and Parasitology, Faculty of Veterinary Medicine, Chattogram Veterinary and Animal Sciences University, Chattogram 4225, Bangladesh

**Author notes:** **Corresponding author:** Yusuke Matsumoto, Transboundary Animal Diseases Research Center Joint Faculty of Veterinary Medicine, Kagoshima University, 1-21-24 Korimoto, Kagoshima 890-0065, Japan, E-mail.

## Abstract

In most paramyxoviruses, RNA editing in the P gene enables expression of the V protein. Human parainfluenza virus type 1 (HPIV-1) differs from most paramyxoviruses in that it lacks RNA editing and does not produce a functional V protein, although its genome retains sequences corresponding to the ancestral V reading frame. Here, we analyzed all HPIV-1 genome sequences available in the NCBI GenBank database to assess the evolutionary state of this V protein–specific region. Using Sendai virus (SeV) as a closely related reference with an identical P gene length, we defined a pseudo-V reading frame by virtually inserting a single nucleotide at the conserved RNA editing site. In this pseudo-V frame, HPIV-1 showed a marked excess of stop codons within the 253-amino-acid region corresponding to the post-editing sequence, far exceeding expectations under random codon usage. This pattern was not observed in other viral genes analyzed under the same definition, nor in SeV, nor was it reproduced by *in silico* evolutionary simulations under constraints preserving the primary open reading frame. These results are consistent with a virus-specific evolutionary trajectory following the loss of RNA editing, rather than with generic coding constraints acting on overlapping reading frames.

## Introduction

In paramyxoviruses that generate multiple gene products through RNA editing, overlapping gene architectures—where a single nucleotide sequence encodes multiple functions—are evolutionarily maintained (1,2). Such structures have been understood as an efficient strategy for maximizing genetic information within a limited genome size (3). However, when one of the functions encoded by an overlapping gene structure becomes dispensable during evolution, how the corresponding genomic region changes and becomes fixed has rarely been examined in a systematic manner.

In most paramyxoviruses, RNA editing occurs in the P gene, in which one or more guanine (G) nucleotides are inserted at a defined probability during transcription (2,4–6). In respiroviruses, including Sendai virus (SeV), insertion of a single G nucleotide predominates (7). This insertion causes a frameshift, resulting in expression of the V protein, a gene product distinct from the P protein. While the region upstream of the editing site is shared between the P and V proteins, the sequence downstream of the insertion constitutes a V protein–specific region encoded in an alternative reading frame (8). In contrast, some paramyxoviruses lack RNA editing altogether. Human parainfluenza virus type 1 (HPIV-1) and Cedar virus (CedV) have lost the RNA editing signal within the P gene, and RNA-edited mRNAs are not detected in infected cells (9–11). Despite the loss of RNA editing, how the genomic region corresponding to the ancestral V protein has evolved, and under what constraints it undergoes pseudogenization, remain poorly understood.

HPIV-1 is phylogenetically closely related to SeV, and the P genes of the two viruses are identical in length, comprising 1,707 nucleotides. This complete length identity allows the RNA editing site to be defined at precisely the same coordinate in both viruses, making it possible to compare them under conditions where the presence or absence of RNA editing is the sole variable. This property makes HPIV-1 and SeV an exceptionally informative comparative system. In SeV, RNA editing occurs immediately after nucleotide position 949 of the P gene, where insertion of a single G nucleotide generates a V-specific reading frame. In this frame, the first stop codon appears at the 69th codon downstream of the insertion site, defining a functional V-specific region.

In the present study, we defined a corresponding reading frame in HPIV-1 by virtually inserting a single nucleotide at the same position, and refer to this frame as the pseudo-V reading frame. In SeV, the region from the insertion site to the first stop codon spans 253 amino acids and contains 12 stop codons in total (Fig, 1A and B). In contrast, in an HPIV-1 representative strain (Acc. No. PV660326.1), the pseudo-V region of the same length contains 29 stop codons, suggesting a pronounced enrichment of stop codons in the sequence corresponding to the V-specific region following the loss of RNA editing (Fig. 1 C and D). Here, we performed a comprehensive analysis of all HPIV-1 genome sequences available in the GenBank database at the National Center for Biotechnology Information (NCBI), examining the positions and frequencies of stop codons within the pseudo-V reading frame. Through this analysis, we aimed to elucidate how overlapping gene architectures are evolutionarily maintained in paramyxoviruses lacking RNA editing, and how one of the encoded functions is progressively lost under virus-specific evolutionary constraints.

**Figure 1.**
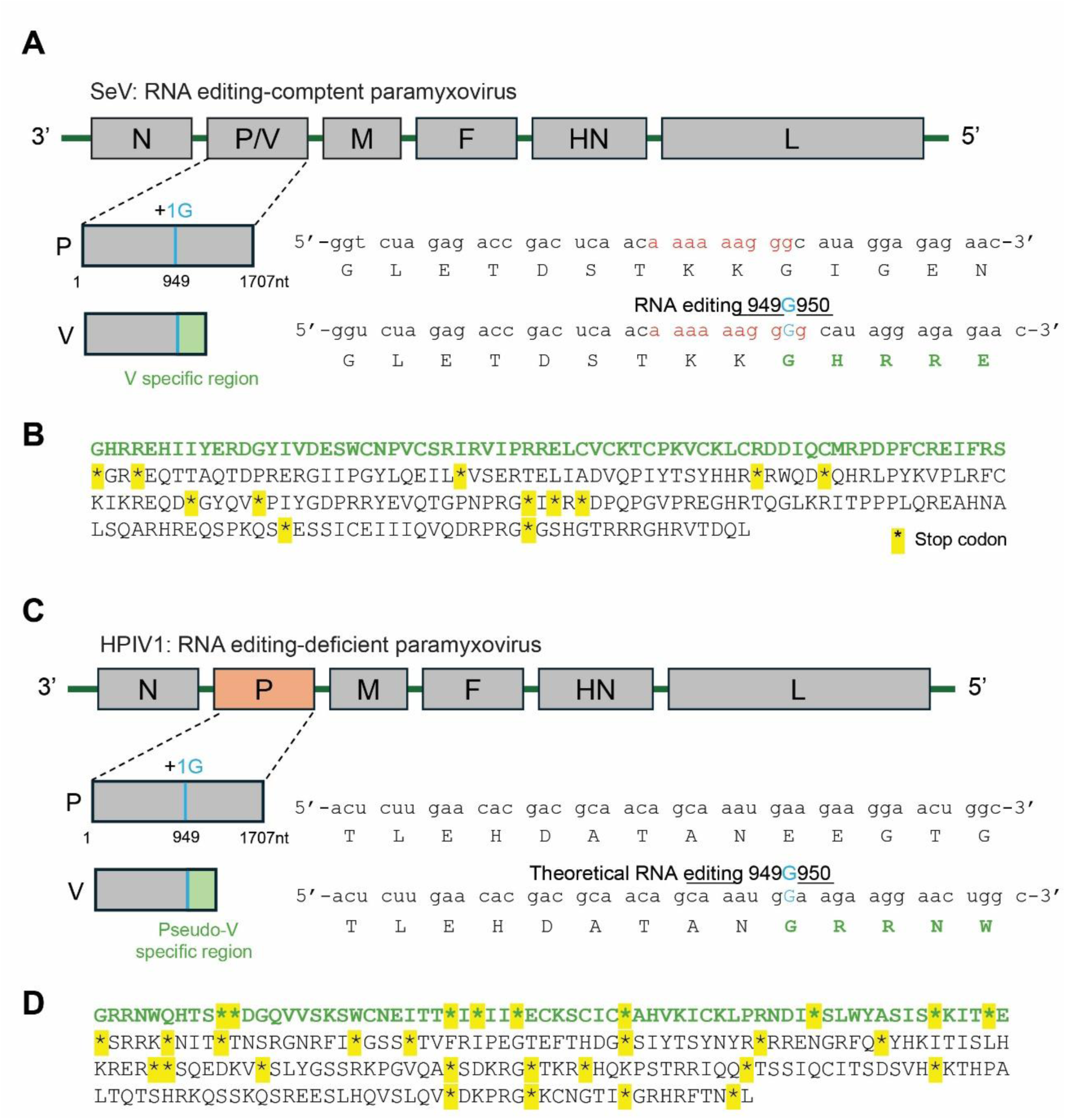
RNA editing and V protein expression in SeV and HPIV-1. **(A)** Schematic representation of the SeV genome, highlighting the P gene and the V gene. The nucleotide sequence of the P gene and its corresponding amino acid translation are shown. RNA editing occurs between nucleotide positions 949 and 950 relative to the 5′ ATG initiation codon of the P gene, resulting in a +1 frameshift. **(B)** Amino acid sequence corresponding to the 253-aa region of the SeV V reading frame generated by RNA editing, showing the distribution of stop codons. The functionally defined V-specific region (from the editing site to the first stop codon) is indicated in bold green. **(C)** Schematic representation of the HPIV-1 genome, highlighting the P gene and the pseudo-V reading frame. The pseudo-V reading frame was defined by theoretical insertion of a single G nucleotide between positions 949 and 950 relative to the 5′ ATG initiation codon of the P gene. **(D)** Amino acid sequence corresponding to the 253-aa pseudo-V region of HPIV-1, translated from the reading frame generated by the virtual nucleotide insertion shown in (C), with stop codons indicated. The region corresponding to the functionally defined V-specific region in SeV (from the editing site to the first stop codon) is indicated in bold green, and the same length was applied to HPIV-1 to enable direct comparison.

## Material and methods

### Sequence selection and definition of the pseudo-V reading frame in HPIV-1

Complete genome sequences of *human respirovirus 1* were retrieved from the NCBI GenBank database. In total, 323 database entries annotated as P gene or phosphoprotein gene were identified. These entries were derived from 243 complete HPIV-1 genome sequences. To enable analyses involving virtual nucleotide insertion at the RNA editing site, corresponding to nucleotide position 949 of the P gene, sequences shorter than 949 nucleotides were excluded, thereby removing entries registered solely as the C protein reading frame. As a result, 243 sequences were retained. Among these 243 sequences, 240 had an identical P gene length of 1,707 nucleotides. Only these 240 sequences were used for subsequent analyses to ensure consistency in reading frame definition and positional correspondence across strains. For each of the 240 P gene sequences, a pseudo-V reading frame was defined by virtual insertion of a single G nucleotide immediately downstream of nucleotide position 949 of the P gene. Following this insertion, the downstream +1 frameshifted reading frame was extracted and treated as the pseudo-V reading frame. The positions of stop codons within this pseudo-V reading frame were identified for each strain.

### Stop codon counting in pseudo-translated reading frames of the N, P, M, F, and HN genes of HPIV-1 and SeV

To quantify stop codon occurrence in the pseudo-V reading frame of the HPIV-1 P gene, the total number of stop codons was calculated for each strain. From each pseudo-V reading frame defined as described above, a 253–amino-acid region downstream of the insertion site was extracted, and all stop codons within this region were counted. To assess whether stop codon accumulation was specific to the P gene, the same procedure was applied to the N, M, F, and HN genes of HPIV-1. For each gene, a control pseudo-translated reading frame was generated by inserting a single G nucleotide immediately upstream of the nucleotide located 758 bases from the 3′ end of the gene, thereby introducing a one-nucleotide frameshift. From each frameshifted sequence, a 253–amino-acid region was extracted, and the total number of stop codons was determined. The L gene was excluded from this analysis because of its large size, as variation in the choice of the pseudo-translated region would substantially influence the results.

The same stop codon counting procedure was also applied to SeV. For each available SeV strain, pseudo-translated reading frames were generated for the P, N, M, F, and HN genes using the same relative insertion position (758 bases from the 3′ end), and stop codons within the corresponding 253–amino-acid regions were counted in an identical manner.

### Forward simulation of overlapping reading frames

To examine whether an increased frequency of stop codons in a pseudo-reading frame can arise as a passive consequence of coding constraints imposed on a main open reading frame (ORF), we performed forward evolutionary simulations under a minimal model. An ancestral nucleotide sequence of length 759 nt was generated randomly by sampling nucleotides (A, C, G, and T) with equal probability. The only constraint imposed on the ancestral sequence was the complete absence of stop codons (TAA, TAG, and TGA) in the main reading frame (frame 0). No other sequence features, such as codon usage bias, GC content bias, or virus-specific motifs, were introduced. Two populations, each consisting of 300 identical copies of the ancestral sequence, were initialized. One population evolved under selection acting exclusively on the main reading frame (“selected population”), whereas the other population evolved neutrally (“neutral population”). Evolution proceeded in discrete generations. In each generation, point mutations were introduced independently at each nucleotide position with a per-site mutation probability of 2 × 10⁻⁴. Insertions and deletions were not allowed. In the selected population, sequences that acquired at least one stop codon in the main reading frame after mutation were discarded, and the next generation was formed by random sampling with replacement from the remaining sequences. If no mutated sequence satisfied the constraint in a given generation, the parental population was retained. In the neutral population, all mutated sequences were propagated without selection, and the next generation was formed by random sampling with replacement. Each simulation was run for 2000 generations, a duration sufficient for the statistics of stop codon usage in the overlapping frame to approach a stationary regime under the imposed constraint. After the final generation, the frequency of stop codons in the +1 reading frame (pseudo-V frame) was calculated by pooling all sequences within each population and counting stop codons relative to the total number of codons in that frame. To account for stochastic variation inherent to finite populations, the simulation was repeated 10 times using different random seeds, with each run representing an independent evolutionary trajectory under identical parameter settings.

## Results

### Stop codons occur at nearly identical positions among HPIV-1 strains in the pseudo-V reading frame

Complete genome sequences of human respirovirus 1 (HPIV-1) were retrieved from the NCBI GenBank database as of December 24, 2025. A total of 323 database entries annotated as P gene or phosphoprotein gene were identified, derived from 243 complete HPIV-1 genome sequences. To perform analyses involving virtual insertion of a single nucleotide downstream of the RNA editing site, corresponding to nucleotide position 949 of the P gene, sequences shorter than 949 nucleotides were excluded, thereby removing entries registered solely as the C protein reading frame (12). As a result, 243 sequences were retained. Of these 243 sequences, 240 had an identical P gene length of 1,707 nucleotides. These 240 sequences were therefore used for subsequent analyses. For each sequence, a single G nucleotide was virtually inserted immediately downstream of nucleotide position 949 of the P gene, and the positions of stop codons in the resulting downstream pseudo-V reading frame were examined across the entire remaining length of the gene. The full downstream region was analyzed to allow a consistent quantitative comparison of stop codon frequency and positional conservation independent of the location of the first stop codon. Figure 2 shows the distribution of stop codon positions within the pseudo-V–specific region, indicating for each position the percentage of strains (out of 240) in which a stop codon was present. Notably, 20 positions were identified at which stop codons were present in more than 95% of the strains (Fig. 2). These results demonstrate that, in the pseudo-V reading frame of HPIV-1, stop codons occur at highly conserved positions across strains.

**Figure 2.**
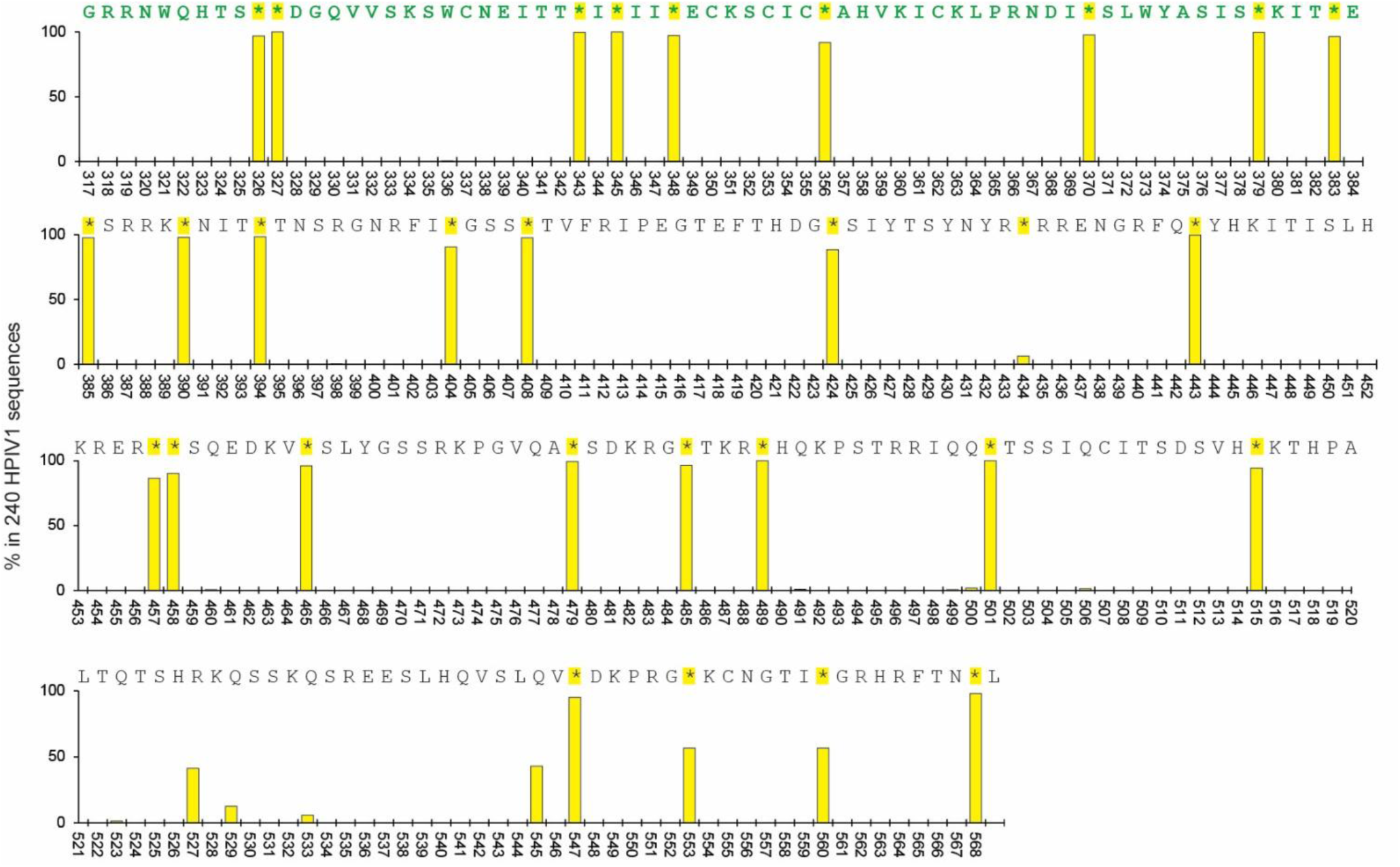
Positions of stop codons within the pseudo-V reading frame of HPIV-1. The 253-aa sequence translated from the HPIV-1 pseudo-V reading frame generated by virtual insertion of a single nucleotide at the RNA editing position is shown. Bar plots indicate the positional frequency of stop codons across 240 HPIV-1 strains. For each amino acid position in the pseudo-V reading frame, the bar height represents the percentage of strains in which a stop codon was present at that position. A value of 100% indicates that all 240 strains contained a stop codon at the corresponding position.

### Stop codons accumulate at a high and non-random frequency in the pseudo-V reading frame of the HPIV-1 P gene

Next, we quantified the total number of stop codons in the pseudo-V–specific region of the HPIV-1 P gene for each strain. The pseudo-V region was defined as the reading frame generated by virtual insertion of a single G nucleotide immediately downstream of nucleotide position 949 of the P gene. This position corresponds to a point located 758 nucleotides upstream of the 3′ end of the gene. To determine whether stop codon accumulation was specific to the P gene, we performed the same analysis on the HPIV1 N, M, F, and HN genes. For each gene, a control pseudo-translated reading frame was generated by inserting a single G nucleotide immediately upstream of the nucleotide located 758 bases from the 3′ end, thereby creating a one-nucleotide frameshift. The downstream length analyzed for each gene was defined to match the length of the region analyzed in the P gene following virtual insertion at the editing-site-equivalent position, enabling direct comparison under an identical analytical framework. The number of stop codons within a 253–amino-acid region was then calculated for each gene. The L gene was excluded from this analysis because of its large size (13), which makes selection of a comparable downstream region inherently ambiguous.

Because 3 of the 64 possible codons are stop codons, the expected probability of stop codon occurrence under random codon usage is 3/64 (4.69%). Based on this probability, the expected number of stop codons in a 253–amino-acid region is 11.86. In the HPIV1 N, F, and HN genes, 13–22 stop codons were observed, corresponding to values comparable to or slightly higher than the expectation (Fig. 3A). In contrast, the M gene contained 7–12 stop codons, showing values comparable to or slightly lower than the expected number (Fig. 3A). Strikingly, the pseudo-V reading frame of the P gene contained 23–30 stop codons, a markedly higher number than observed in any of the other genes (Fig. 3A). The same analysis was performed for SeV (Fig. 3B). Analysis of 20 available SeV strains revealed 11–21 stop codons in all genes examined, and no gene-specific enrichment of stop codons was observed in the SeV pseudo-V reading frame of the P gene (Fig. 3B). These results indicate that, in HPIV-1, an unusually high frequency of stop codons is specifically confined to the pseudo-V region of the P gene.

**Figure 3.**
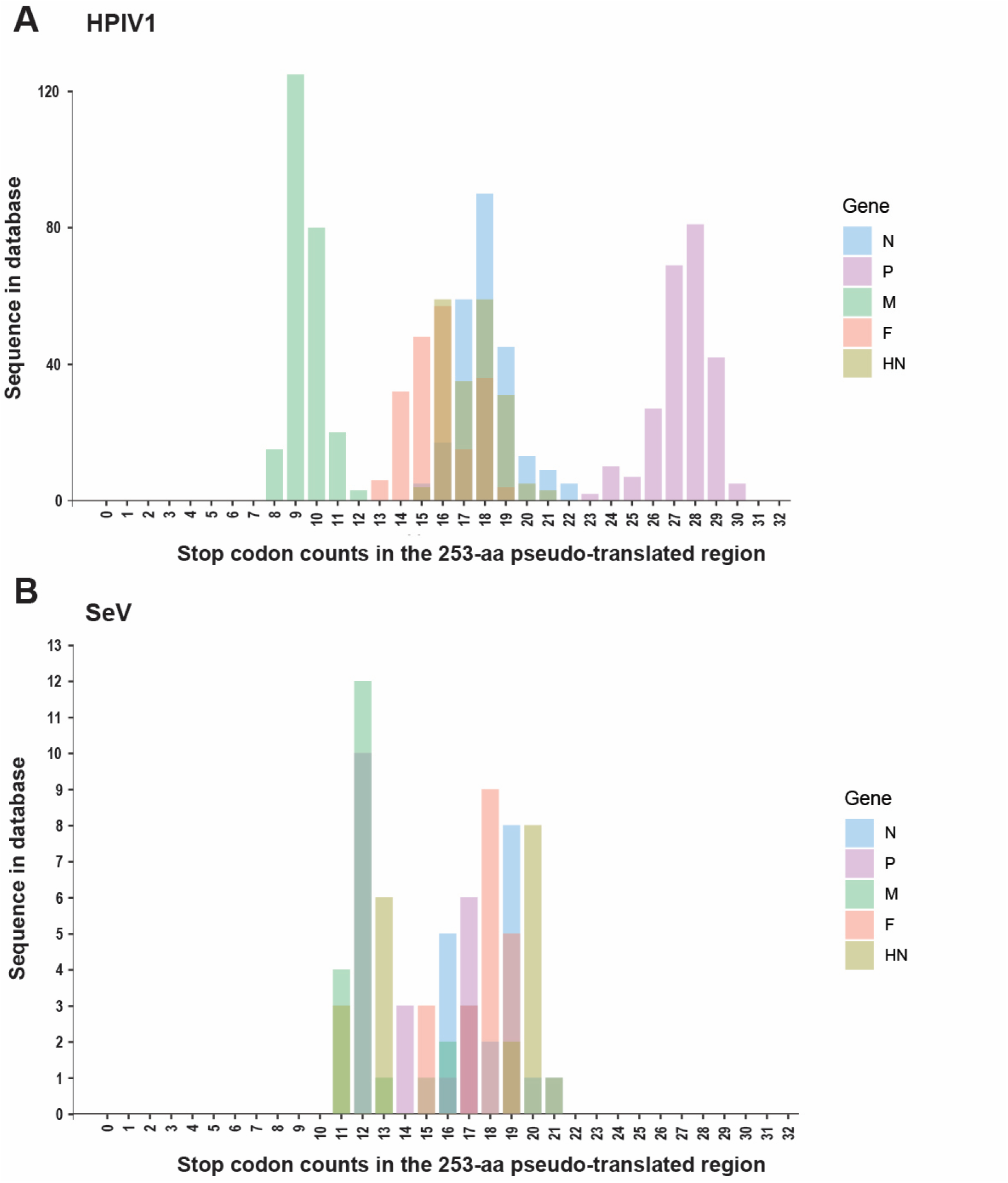
Frequency of stop codons in the in the 253-aa pseudo-translated region. **(A)** Stop codons were counted in a 253-aa pseudo-translated region generated by virtual insertion of a single G nucleotide immediately downstream of nucleotide position 949 of the P gene (equivalent to insertion immediately upstream of the nucleotide located 758 bases from the 3′ end of each gene) in all 240 HPIV-1 sequences available in the database. The analysis was performed separately for the N, P, M, F, and HN genes. The y-axis indicates the number of sequences in the database showing a given stop codon count. **(B)** The same analysis was performed for SeV using 20 sequences available in the database.

For the 240 HPIV-1 strains analyzed, the number of stop codons per strain was examined in relation to year of isolation, geographic origin, and phylogenetic relationships inferred from P gene sequences (Fig. 4). Although a small number of strains isolated at earlier time points showed slightly fewer stop codons, no clear trend of progressive accumulation of stop codons over time was observed. In addition, no geographic bias in stop codon frequency was detected.

**Figure 4.**
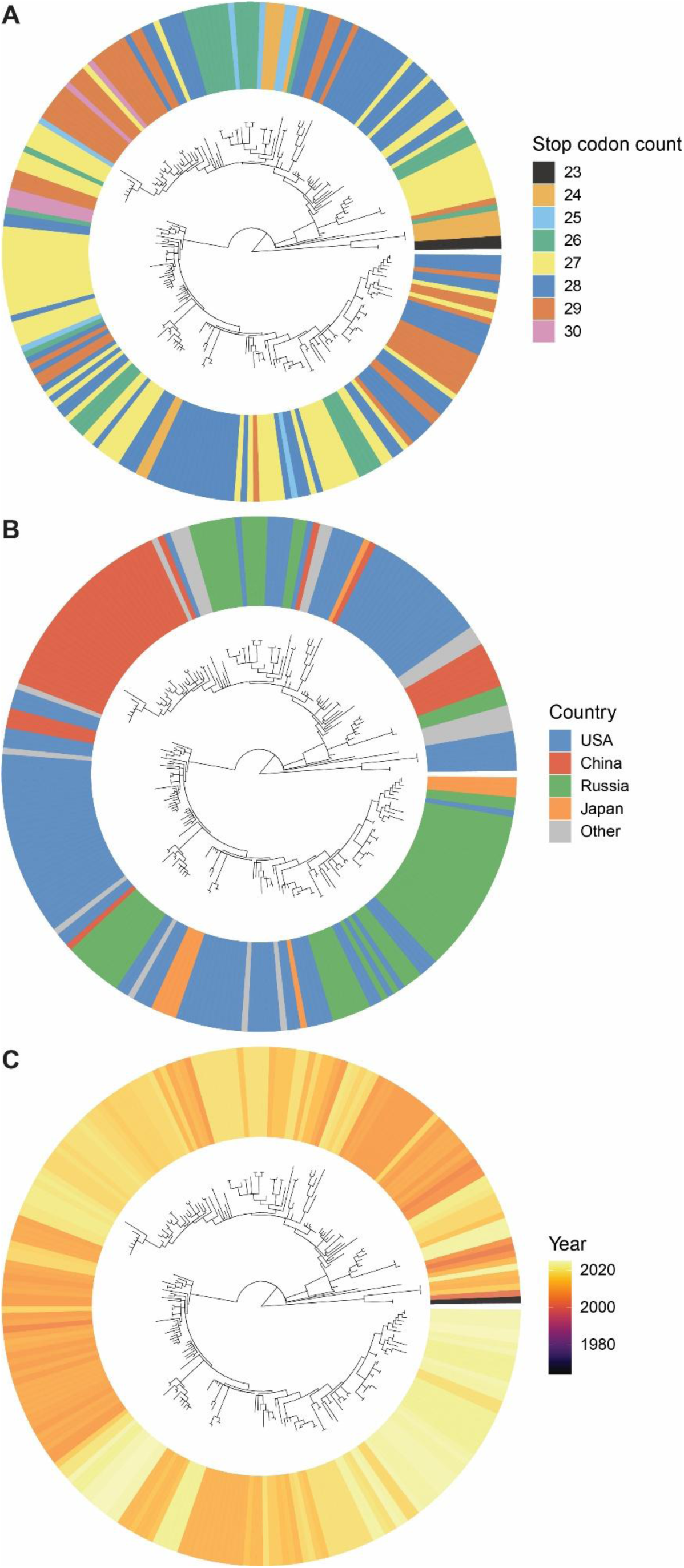
Relationship between stop codon counts in the pseudo-V region of the HPIV-1 P gene and strain metadata. **(A)** Circular phylogenetic tree constructed from the P gene sequences of 240 HPIV-1 strains. An outer circular ring indicates the number of stop codons (ranging from 23 to 30) present in the 253-aa pseudo-V region of each strain, with colors representing different stop codon counts. **(B)** The same phylogenetic tree as in (A), with the outer ring colored according to the country of isolation of each strain. **(C)** The same phylogenetic tree as in (A), with the outer ring colored according to the year of isolation or year of sequence registration for each strain.

### The stop codon accumulation observed in HPIV-1 could not be explained solely by coding constraints that exclude stop codons from the primary ORF

In the pseudo-V reading frame of the HPIV-1 P gene, stop codons were present at high frequency. We therefore tested, using *in silico* evolutionary simulations, whether this pattern could be reproduced by imposing only a constraint that excludes stop codons from the primary ORF.

In these simulations, an ancestral sequence of 759 nucleotides was generated at random, under the sole constraint that the primary ORF (frame 0) contained no stop codons. A population consisting of 300 identical copies of this ancestral sequence was initialized, and point mutations were introduced independently at each nucleotide position in each generation (per-nucleotide mutation probability, 2 × 10⁻⁴). Insertions and deletions were not permitted. In one population, sequences that acquired stop codons in the primary ORF after mutation were removed by selection (selected population), whereas in the other population, sequences were allowed to evolve neutrally without selection (neutral population). Each simulation was run for 2,000 generations, and 10 independent replicates were performed using different random seeds.

At the final generation of each replicate, all sequences in the population were pooled, and the frequency of stop codons in the +1 reading frame (pseudo-V frame) was calculated. In the selected population, stop codon frequencies ranged from 3.3 to 6.7%, which was comparable to the range observed in the neutral population (3.5–6.5%). Among the 10 independent replicates, stop codon frequency in the selected population was higher than that in the neutral population in six replicates and lower in four replicates, indicating no consistent directional difference between the two conditions.

Taken together, these results indicate that, under a model assuming only point mutations and a coding constraint that strictly excludes stop codons from the primary ORF, no consistent increase in stop codon frequency is reproduced in the overlapping pseudo-V reading frame. Thus, accumulation of stop codons in the pseudo-V region cannot be explained solely by coding constraints acting on the primary ORF.

## Discussion

In this study, we found that the pseudo-V reading frame of the HPIV-1 P gene contains a substantially higher number of stop codons than any other gene analyzed under the same definition, a pattern not observed in the HPIV-1 N, M, F, or HN genes or in the closely related SeV. These results indicate that the elevated stop codon frequency observed in this region is not readily explained solely by general properties of pseud-translated reading frames of by the analytical framework used here.

In viruses with overlapping gene architectures, strong constraints act on the primary ORF to prevent premature translational termination by suppressing the occurrence of stop codons. It is intuitively plausible that such constraints could passively influence stop codon frequencies in alternative reading frames encoded by the same nucleotide sequence. However, forward *in silico* evolutionary simulations showed that, under a model incorporating only point mutations and a constraint excluding stop codons from the primary ORF, no consistent directional increase in the total number of stop codons was reproduced in the pseudo-V reading frame (Fig. 5). These results indicate that the stop codon accumulation observed in the pseudo-V frame of HPIV-1 is not a computationally inevitable consequence of overlapping reading frames.

**Figure 5.**
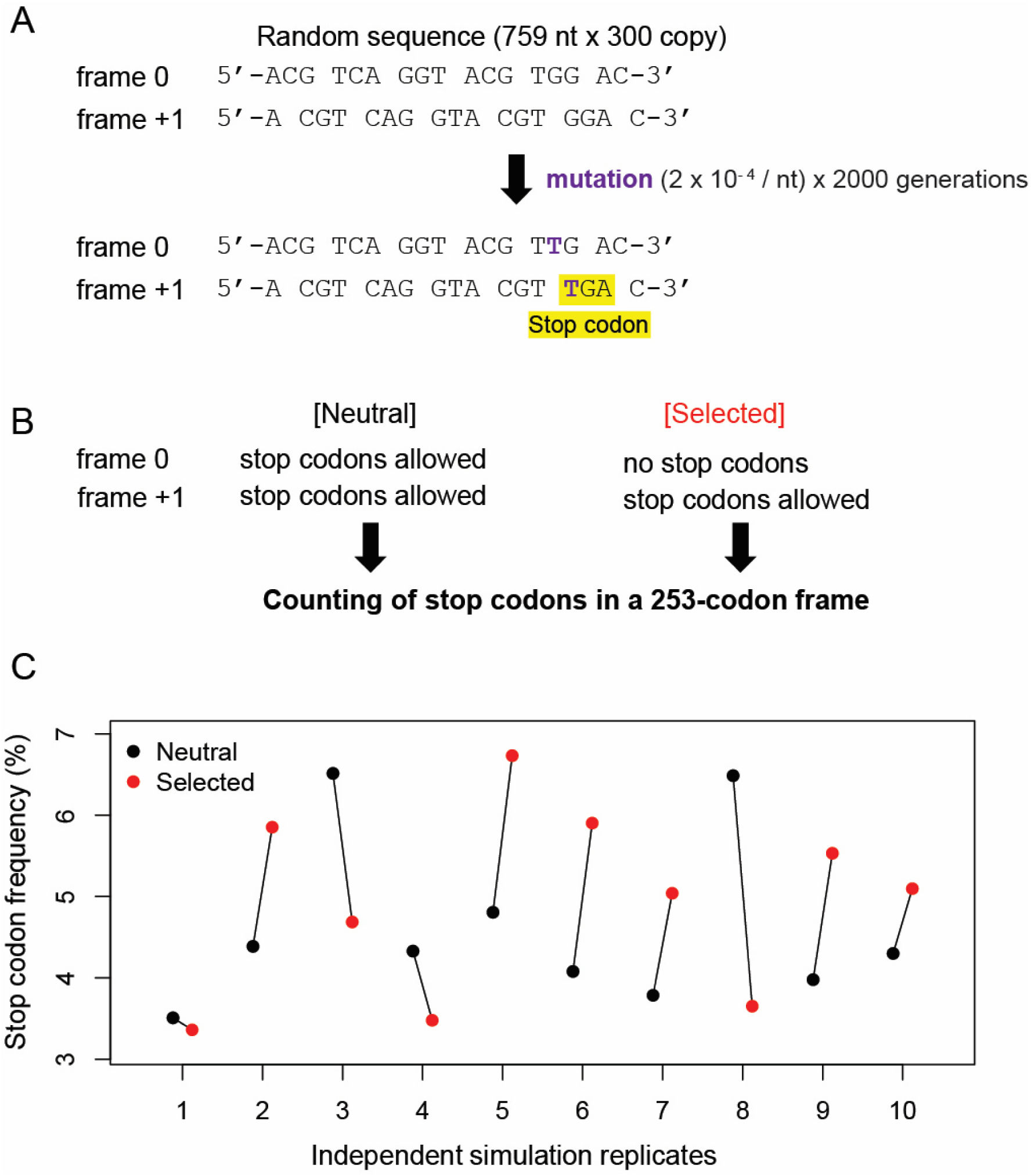
In silico evolutionary simulations of stop codon accumulation in a +1 frameshifted reading frame. **(A)** Schematic overview of the simulation framework. An ancestral nucleotide sequence of 759 nt was generated at random and used to initialize a population of 300 identical copies. Point mutations were introduced independently at each nucleotide position with a mutation rate of 2 × 10⁻⁴ per nucleotide per generation over 2,000 generations. No insertions or deletions were allowed. The primary reading frame (frame 0) and the overlapping +1 frameshifted reading frame are shown, illustrating the appearance of stop codons in the +1 frame during evolution. **(B)** Definition of selection regimes. In the neutral condition, stop codons were permitted in both frame 0 and the +1 frame. In the selected condition, sequences containing stop codons in frame 0 were eliminated, whereas stop codons were allowed in the +1 frameshifted reading frame. After evolution, stop codons were counted within a 253-codon window in the +1 reading frame. **(C)** Stop codon frequency (%) in the +1 frameshifted reading frame for neutral and selected populations across 10 independent simulation replicates. Each point represents the stop codon frequency calculated from the final generation of a single replicate.

Taken together, the results presented here argue against simple models in which stop codon accumulation in the pseudo-V reading frame arises solely from generic constraints acting on the primary ORF. Instead, the observed pattern defines an evolutionary outcome that cannot be accounted for by such models alone and underscores the lineage-specific nature of genome evolution following the loss of RNA editing.

Paramyxovirus V proteins are multifunctional and play important roles in antagonizing host innate immune responses. However, their presence and functionality vary across paramyxovirus lineages. For example, in HPIV-3 multiple stop codons between the editing site and the conserved zinc-finger domain prevent production of a full-length V protein, and only truncated forms lacking the canonical C-terminal region are detected in infected cells, suggesting pseudogenization of the V-specific coding region (2,14). These observations indicate that V protein function is not uniformly conserved and may be subject to lineage-specific evolutionary constraints. Requirements for V protein function may also vary depending on cellular context. Measles virus variants deficient in V expression are positively selected in lymphocytic cells but rapidly outcompeted by V-competent viruses in epithelial cells, demonstrating environment-dependent selection on V protein function (15). Within this broader evolutionary context, the lineage-specific accumulation of stop codons documented here in the pseudo-V reading frame of HPIV-1 likely reflects long-term relaxation of functional constraints following the loss of RNA editing and V protein expression. Although the present study focuses on sequence-level evolutionary outcomes rather than functional consequences, the conserved and non-random patterns identified here provide a framework for future studies examining how loss of V protein function influences genome evolution in paramyxoviruses.

## Data availability

All databases used in this study are available from DDBJ/ENA/GenBank (https://www.ddbj.nig.ac.jp/about/insdc-e.html). The accession numbers of the viral sequences used in this study are listed in Supplementary Table 1.

Sequence selection, definition of pseudo-translated reading frames, and stop codon counting analyses were performed using custom scripts written in R. Forward evolutionary simulations were performed using Python scripts. All scripts used in this study are publicly available on GitHub (https://github.com/ymatsu51/hpiv1-stop-codon-analysis).

